# DBlink: Dynamic localization microscopy in super spatiotemporal resolution via deep learning

**DOI:** 10.1101/2022.07.01.498428

**Authors:** Alon Saguy, Onit Alalouf, Nadav Opatovski, Soohyen Jang, Mike Heilemann, Yoav Shechtman

## Abstract

Single molecule localization microscopy (SMLM) has revolutionized biological imaging, improving the spatial resolution of traditional microscopes by an order of magnitude. However, SMLM techniques depend on accumulation of many localizations over thousands of recorded frames to yield a single super-resolved image, which is time consuming. Hence, the capability of SMLM to observe dynamics has always been limited. Typically, a few minutes of data acquisition are needed to reconstruct a single super-resolved frame. In this work, we present DBlink, a novel deep-learning-based algorithm for super spatiotemporal resolution reconstruction from SMLM data. The input to DBlink is a recorded video of single molecule localization microscopy data and the output is a super spatiotemporal resolution video reconstruction. We use bi-directional long short term memory (LSTM) network architecture, designed for capturing long term dependencies between different input frames. We demonstrate DBlink performance on simulated data of random filaments and mitochondria-like structures, on experimental SMLM data in controlled motion conditions, and finally on live cell dynamic SMLM. Our neural network based spatiotemporal interpolation method constitutes a significant advance in super-resolution imaging of dynamic processes in live cells.

## Introduction

The spatial resolution in optical microscopes is bounded by the diffraction limit, which sets the minimal achievable resolution of standard light microscopes at about half the wavelength of light, corresponding, in the visible range, to ∼200-300 nm. To overcome this limitation, and enable higher resolution, super-resolution microscopy (SRM) methods have been developed. Notable methods of this family include stimulated emission depletion (STED)^1^, structured-illumination microscopy (SIM)^2^, as well as single molecule localization microscopy (SMLM)^3–5^. Prominent variants of SMLM include photoactivated localization microscopy^3^ (PALM), stochastic optical reconstruction microscopy^3^ (STORM), points accumulation for imaging in nanoscale topography^6^ (PAINT), and DNA-PAINT^5^. The SMLM variants differ in their experimental conditions, however they share a similar overall pipeline: First, fluorescent molecules are used to label structures in a specimen. Then, a sequence of frames is captured, in which only a sparse, random subset of molecules emit light per-frame. Subsequently, each emission event is detected, and fit to a model of the system point spread function (PSF) allowing highly precise determination of the emitting fluorophore’s position. Finally, by accumulating the localizations of thousands of emitters, the output of SMLM is a single super-resolved image of the structure, typically with an order of magnitude resolution improvement compared to the diffraction limit.

An inherent limitation in SMLM is its temporal resolution. Accumulating a large enough number (typically millions) of single-molecule emission events, to generate a continuous image takes a long time. Indeed, the typical temporal resolution in SMLM is on the order of minutes, while tens-of-seconds resolutions have also been reported^7,8^. Recent advances in deep learning algorithms have yielded computational algorithms that further improve the capabilities of SMLM. ANNA-PALM^7^ significantly reduces the number of frames needed for super-resolution reconstruction. Deep-STORM^9,10^ as well as DECODE^11^ enable researchers to analyze densely labeled SMLM experiments by training a neural network to perform multi-emitter fitting in super-resolution. Importantly, while these algorithms and other non-SMLM methods^12,13^ perform exceptionally well in visualization of nanoscale structures, they are still mainly applicable for the analysis of static data or very slow dynamics.

The main reason why previous methods are useful mostly for static data is because the typical SMLM reconstruction process does not exploit structural-correlations over long periods of time (longer than the temporal window being reconstructed). On the other hand, algorithms^13,14^ that do exploit temporal interpolation along the video frames, do not apply spatial interpolation per frame; an algorithm that combines both spatial and long-range temporal interpolation would be optimal.

In this paper, we present DBlink, a novel algorithm that increases the spatiotemporal resolution in the reconstruction of dynamic SMLM data. We use a bi-directional long short term memory (LSTM) network, that receives as input a video of super-resolved localization maps and outputs a video of a dynamic super-resolved structure (Fig. 1). The super-resolved localization maps can be obtained by using existing algorithms, e.g. Deep-STORM^15^ or ThunderSTORM^16^. In order to perform such spatiotemporal interpolation, DBlink relies on inter-frame correlations, and on prior information regarding the imaged sample – namely, its type (e.g. microtubules, mitochondria, etc.). We first demonstrate the ability of DBlink to reconstruct super spatiotemporal resolution videos of the dynamics of simulated filaments. Then, we validate our network performance on experimental data of drifting and rotating microtubule filaments. We present super spatiotemporal resolution reconstructions of microtubule dynamics in live cells, achieving spatial resolution of ∼40 nm and temporal resolution of 300 ms (∼20 frames). Finally, we demonstrate the reconstruction of mitochondrial dynamics from live-cell PAINT data using a non-covalent, weak affinity fluorophore label.

**Figure 1:**
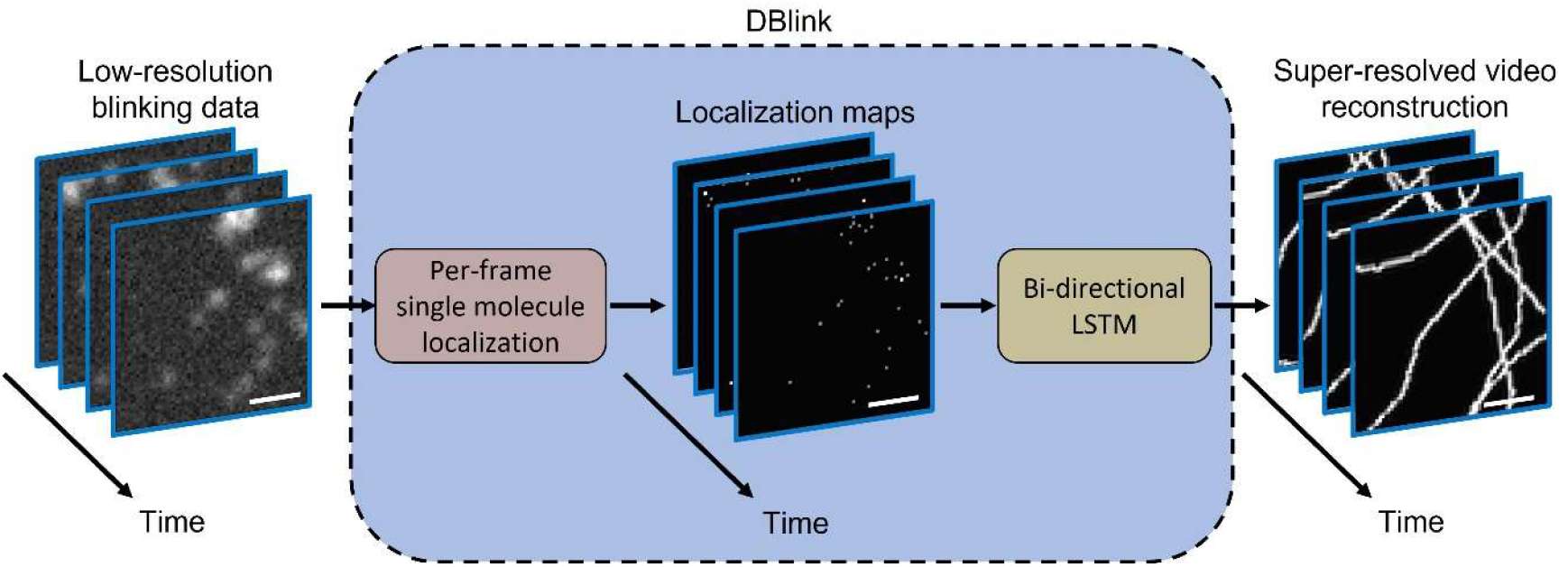
DBlink concept. Low-resolution frames containing stochastic blinking events are analyzed by a localization algorithm, in our case, Deep-STORM^9^, which generates super-resolved localization maps for each input frame. The localization maps serve as input to a LSTM network that provides as output super-resolution video reconstruction of the imaged structure. Scale bar = 2.5 um.

## Results

Our goal is to extend the temporal resolution of SMLM beyond the current state-of-the-art, while maintaining high-spatial resolution. Conceptually, the problem at hand is spatiotemporal interpolation of a dynamic structure from noisy pointwise samples. Clearly, there is insufficient information per frame, and solving this inverse problem requires some sort of prior knowledge. The strategy we chose here is to supply this knowledge in the form of training a neural network on data that resembles the desired structures – spatially as well as temporally. In addition to the exploitation of prior knowledge, we searched for a neural network architecture that can capture long term dependencies between different video frames. LSTM networks have previously proven themselves as a good solution for this task^17,18^. The hidden states of LSTM cells carry information from previous frames as they propagate throughout the video. Since our method analyzes experiment in retrospect, we possess also information from future frames; therefore, we decided to use a bi-directional LSTM network, concatenate the forward pass and the backward pass, and input them into a convolutional neural network (CNN) that provides the final video reconstruction (Supplementary Fig, S1). In the following example applications, we show that this approach is feasible and produces high quality results.

First, we tested our approach on simulated data (Fig. 2). To do so, we generated simulated filaments according to the model of Shariff *et al*.^19^ (see methods section); then, we shifted and rotated them randomly in time (see SI for info). The simulated ground truth for each video was a binary map that contained ones where there was a filament and zeros everywhere else. Then, we generated simulated localizations based on the ground truth mask, at finite precision. To simulate motion within a single acquisition frame, we summed the localizations over temporal windows of 10 simulated-frames. The frame summation window size could be optimized according to different experimental conditions for better performance. The neural network was able to reconstruct random shifting structures over time with high accuracy (Supplementary Video 1), namely, 90% of the simulated binary map matched the predicted structure, and only 1.3% of the predicted structure was hallucinated (see SI for more details). In this simulation, the temporal resolution corresponds to 1 reconstructed frame per 10 simulated blinking frames.

**Figure 2:**
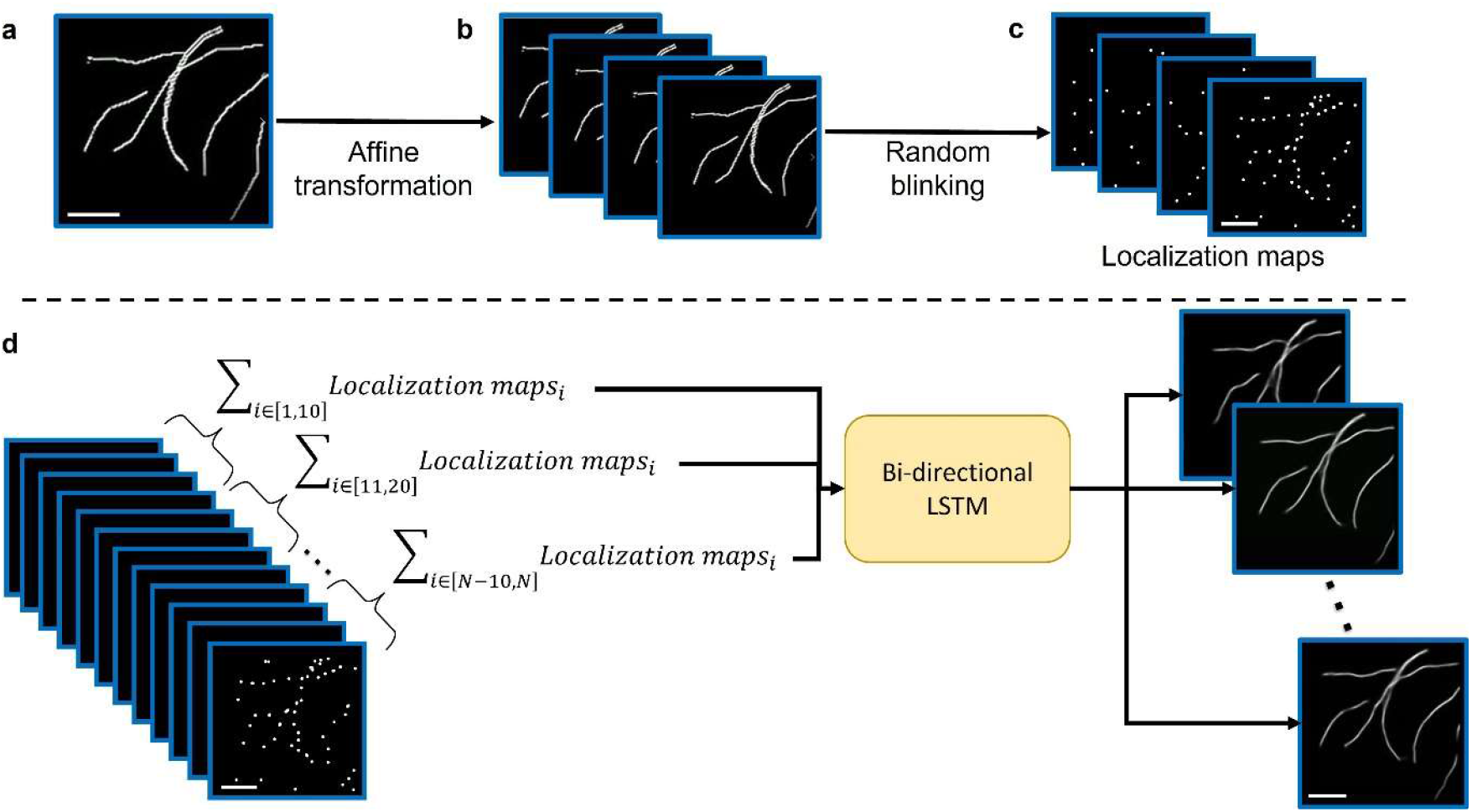
Generation and analysis of simulated filament training data. **A** We simulated a random number of filaments in the field of view (FOV) according to the model of Shariff, et al^19^. **B** Then, we applied gradually increasing affine transformations over a predefined video length of N frames. **c** Next, we generated random blinking event localizations based on the simulated structure. **d** Finally, we summed the simulated localizations every 10 frames and inserted the summed frames (total of N/10 frames) to the LSTM. The output of the LSTM was a super-resolution reconstruction video of length N/10. Scale bar = 2.5 um.

To quantify the spatial reconstruction resolution of the network, we performed Fourier ring correlation analysis^20^ as well as decorrelation analysis^21^ between the network reconstruction and a STORM reconstruction on a static sample. The network result was consistent with standard STORM reconstruction using ThunderSTORM^16^ up to a resolution of ∼37 nm (see SI).

Next, as a first validation of our method on experimental data, we reconstructed a static structure which was shifting laterally over time. For this, we captured a STORM experiment of fixed microtubules exhibiting naturally occurring lateral sample-drift. We estimated the drift using Deep-STORM drift correction mechanism, which is based on cross-correlation, and received a total shift of 240 nm in y direction and 400 nm in x direction (Fig 3). Next, we used the localization maps provided by Deep-STORM and summed them over windows of 100 frames with 50 ms acquisition time, to obtain high enough density per-input-frame to match our network training density. Finally, we input the summed localization maps into our network and received a super spatiotemporal reconstruction of the shifting data (Supplementary Video 2), at a temporal resolution of 0.2 frames per second. We predicted the drift according to the cross-correlation between the first reconstructed frame of our network and every other frame in the reconstructed video. The mean distance between our drift prediction and Deep-STORM prediction over the course of the experiment was 38 nm (Fig. 3). Notably, the network did not have any prior knowledge that the sample is static and drifting, namely, that the only motion was a global shift; rather, the network treated this data as general dynamic data. A global-motion prior would improve the performance significantly, at the cost of a less generalized solution.

**Figure 3:**
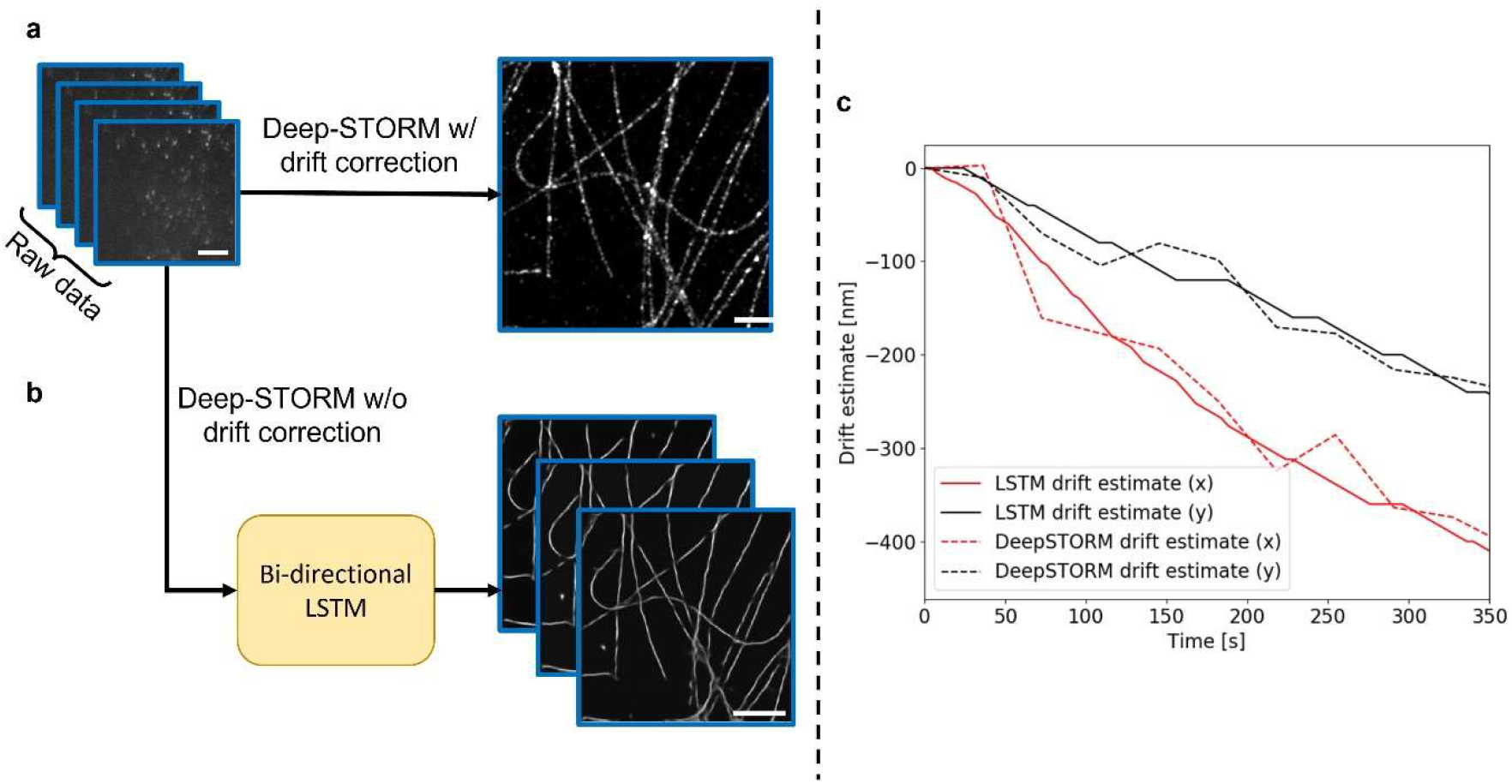
Tracking drifting microtubules. **a** Deep-STORM is used to analyze 10,000 frames of experimental STORM data containing undesirable drift and obtain a super-resolution reconstruction of the microtubule structure. We used Deep-STORM’s drift correction tool to predict the drift over the course of the experiment. **b** The same STORM video is analyzed using DBlink. We have determined the drift by taking the maximal value of the cross-correlation between each reconstructed frame and the initial reconstructed frame. **c** Deep-STORM drift prediction (dashed line) is consistent with the DBlink’s motion prediction. Scale bar = 2.5 *µm*.

To demonstrate our algorithm on a more complex type of motion than lateral shift, while still possessing knowledge of the sample structure to serve as validation, we captured a STORM video of static microtubules while rotating the camera manually (Fig. 4). We added to the sample fluorescent beads to serve as fiducials reporting on sample rotation. The rotation could be predicted by finding the lateral displacement of the bead in each frame and calculating the arctan of y position divided by x position, relative to the rotation axis. At the end of the experiment, we stopped rotating the camera and let the blinking continue for ∼15,000 more frames; this is to generate the ground truth structure by running Deep-STORM on the static portion of the video. To test our reconstruction performance, we computationally rotated back the predicted structure in each frame according to the calculated rotation angle. Then, we compared the computationally-rotated video to the static reconstruction obtained by Deep-STORM (Supplementary Video S3). To quantify the prediction error, we measured the consistency between the reconstructed video frames based on the cross-correlation between every two frames in the reconstructed video, achieving a mean consistency score of 0.91, which indicates a stable reconstruction (see SI). In this experiment, we used a window size of 40 summed frames to generate the reconstructed video, with acquisition time of 20 ms per frame, resulting in temporal resolution of 1.25 frames per seconds.

**Figure 4:**
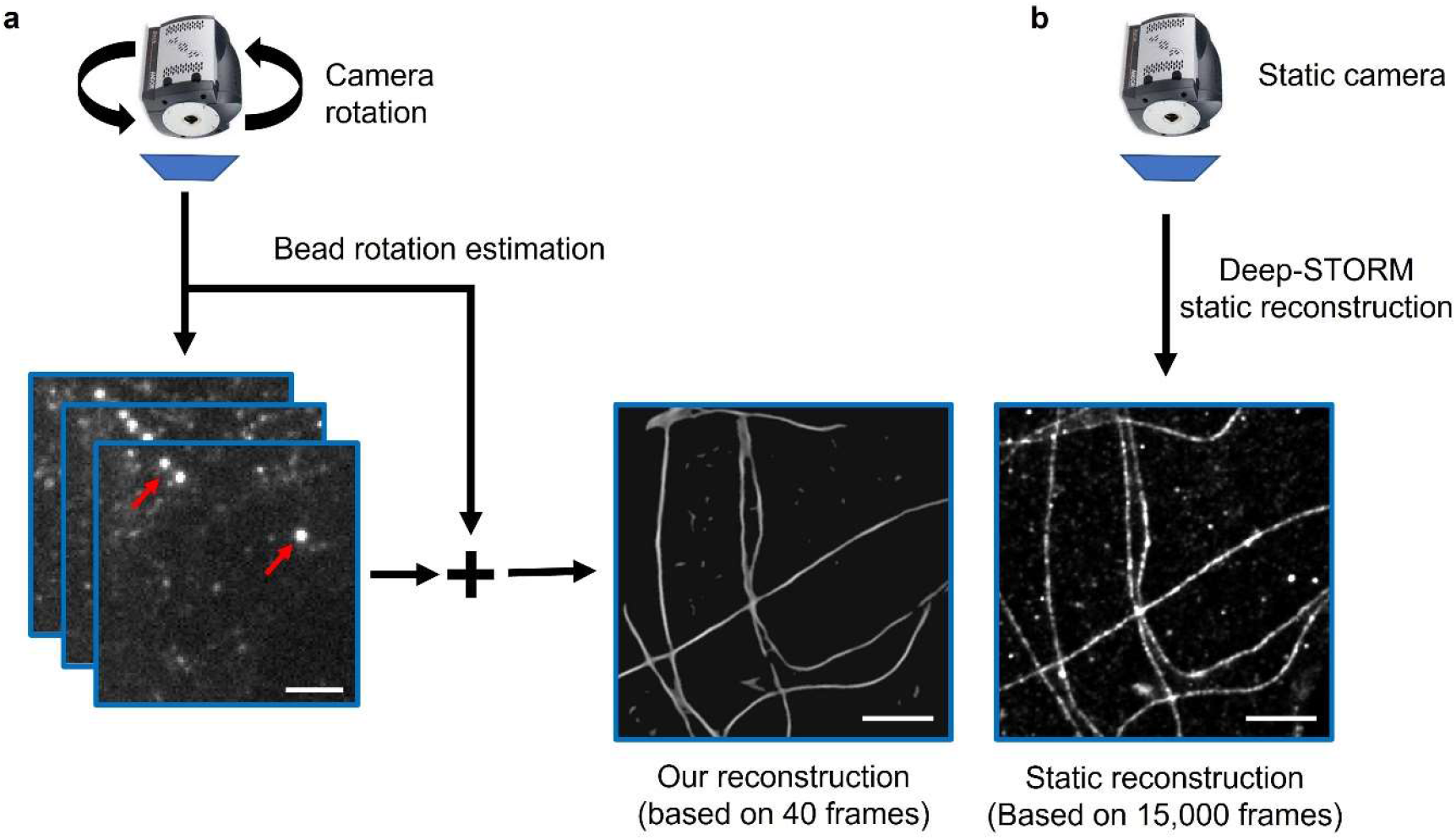
DBlink performance quantification using rotating microtubules. **a** A STORM movie of blinking microtubules is captured while rotating the camera. Fluorescent beads (red arrows) serve as fiducials reporting on the rotation. **b** Rotation is stopped, and a static STORM video is captured for 15,000 more frames, used to produce a ground truth static structure via Deep-STORM. The static structure is then compared to each frame in the dynamic reconstructed video, rotated appropriately. Scale bar = 2.5 *µm*.

Next, we tracked microtubule dynamics in live cells. Since ground truth information was not available in this case, we compared our network reconstructions to two alternative solutions: Deep-STORM reconstructions based on short time windows, and a previously reported blind inpainting algorithm^8^ (Supplementary Video 4), meant to compensate on the loss of information that occurs when choosing short temporal STORM windows. In this experiment, the input to DBlink was the sum of localizations over windows of 20 frames. We set the number of input frames for Deep-STORM windows and blind inpainting to be 300 frames. This number was chosen by optimizing the window size for the best reconstruction result (see Supplementary Fig. S6).

Qualitatively, DBlink reconstructions consistently outperformed the other two methods. Although blind inpainting has managed to filter most of the noise in Deep-STORM data, it performed poorly in densely labeled areas. Furthermore, rapid dynamics caused motion blur in temporally-windowed STORM reconstructions (Fig. 5), while our network provided a more stable reconstruction in areas exhibiting rapid motion. The spatial resolution we achieved was 40 nm and the temporal resolution was 0.3 seconds, namely, 3.3 frames per second.

**Figure 5:**
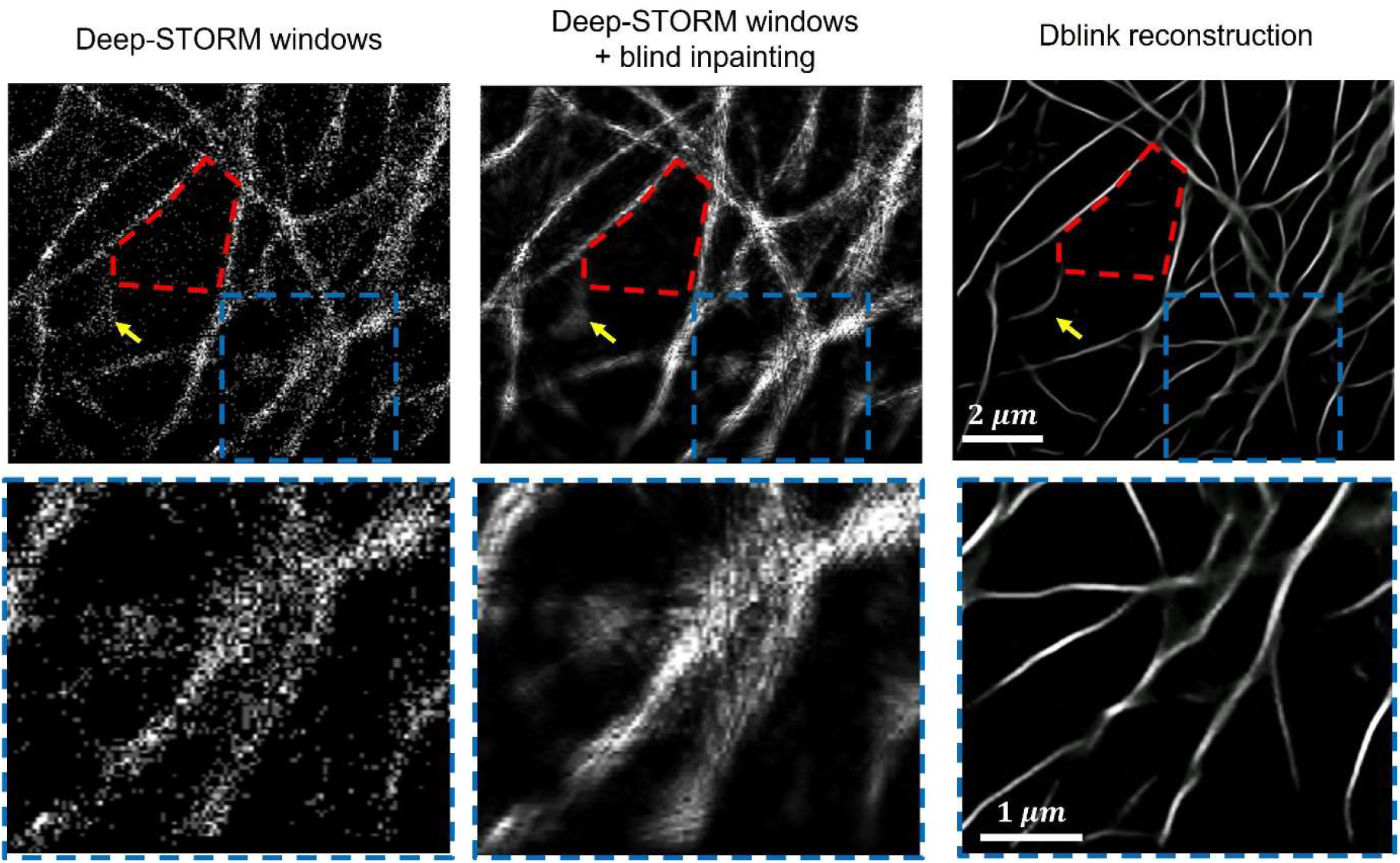
DBlink reconstruction of dynamic microtubule filaments in a live cell. We compared DBlink to state-of-the-art single-molecule super-resolution reconstruction methods. Left column: a single reconstructed frame of microtubule dynamics obtained by summing Deep-STORM localizations over 300 frames. Middle column: reconstruction after summing 300 localization frames and applying blind inpainting. Right column: DBlink reconstruction. Bottom row: a region of interest (blue dashed box) for emphasizing the differences between the different reconstructions. The red dashed polygon marks the noise cancellation performed by both blind inpainting and our algorithm. The yellow arrow marks motion blur artefacts caused by summing localization from multiple frames. Upper row scale bar = 2 *µm*; bottom row scale bar = 1 *µm*.

Finally, we reconstructed the dynamics of mitochondria in live cells from high-density single-molecule data of mitochondrial protein COX8. In addition, we extended the observation time in live cell imaging by using a HaloTag7 fusion in combination with a non-covalent, weak affinity fluorophore tag that binds to and unbinds from the target and acts as an exchangeable fluorophore label^22^ (Supplementary Video 5). In this case, training required a model for mitochondrion size and shape, labeling density, motion type and speed, etc. For this purpose, we developed a dynamic mitochondria simulator (see SI for more information). After training the neural network, we analyzed a SMLM video containing dynamic mitochondria labeled with HaloTag (see Methods section). The mitochondria displayed contraction, elongation, and drift, at different velocities, similarly to previously published work on mitochondrial dynamics^23^. DBlink managed to detect the dynamics with high fidelity to the observed data (Fig. 6); this validates the generalizability and applicability of our network for the analysis of various biological samples, contingent on appropriate training.

**Figure 6:**
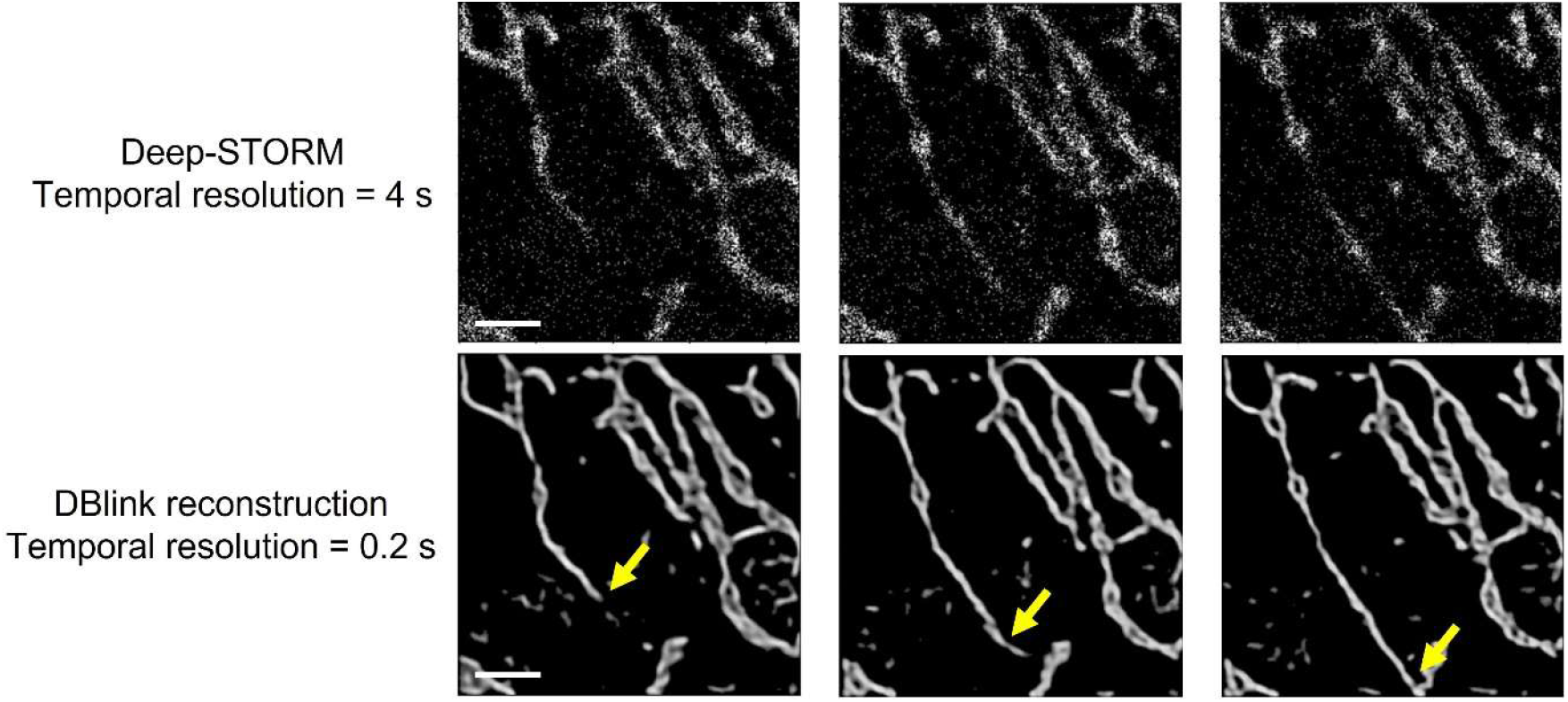
DBlink reconstruction of mitochondria dynamics in a live cell. Upper row: 10-frame summation of Deep-STORM localizations in three different timepoints of the experiment (0, 5, 15 s). Bottom row: DBlink reconstructions of dynamic mitochondria. Yellow arrow marks mitochondria elongating over time. Scale bar = 5 *µm*.

The temporal resolution we achieved in this experiment is 5 frames per second and the spatial resolution is 75 nm, calculated by decorrelation analysis^21^.

## Discussion

In this paper, we presented a method for super spatiotemporal resolution reconstruction of dynamic SMLM data. Our solution utilizes two main assumptions: 1) the imaged-sample class is known (e.g. filaments, mitochondria), and 2) dynamically varying objects maintain some degree of structural similarity over time, allowing the network to exploit structural information correlation over time. In other words – the information used by our network to recover an SMLM video with a temporal rate of 20 frames is not contained in these 20 frames alone – but rather also in a window of hundreds of frames around it. Notably, both assumptions are necessary in order to achieve the spatiotemporal resolutions demonstrated in this work.

To overcome the challenge of verifying the network results, we tested several cases in which estimation of ground truth position was possible, including numerical simulation, sample drift, and controllable whole-sample motion. First, we proved our network can reconstruct random motion of filaments in simulations. The network could reconstruct nanoscale rapid movement of simulated filaments with 90% of the simulated structure correctly classified per simulated video, assuming certain SMLM conditions (e.g. ∼20 nm localization precision, ∼1 emitter per µm^2^); naturally, predication accuracy varies as a function of fluorophore density, motion speed and other experimental parameters (see SI for more details). Next, an experiment that consisted of undesirable sample drift was performed. In this experiment, we could obtain the dynamic structure, since the sample was fixed and only global drift was present, which could be estimated by Deep-STORM. Our drift prediction and the structure reconstruction agreed with the predictions of Deep-STORM (mean error of 38 nm). Additionally, we demonstrated the ability of our network to reconstruct a different global motion: whole sample rotation. These two validation experiments show that our network manages to reconstruct the ground truth structure at high spatiotemporal resolution with high fidelity.

Ultimately, the goal of our method is to enable live SMLM. For this purpose, we have analyzed SMLM videos of live-cell microtubule dynamics, provided by R. Tachibana, et al.^24^. Since no ground truth structure is available in such an experiment, we have qualitatively assessed the reconstruction accuracy by comparison to another state-of-the-art solution for dynamic SMLM. We used Deep-STORM to generate localizations over temporal windows, followed by blind inpainting^8^ to filter noisy localization and compensate for missing information. Both blind inpainting and our algorithm have successfully managed to filter noisy localizations and compensate on missing information. Nevertheless, our network was able to reliably reconstruct much denser areas in the FOV and without motion blur, which was possible since each input contained localizations from only 20 recorded frames. Temporally-windowed STORM over short windows performs poorly because information regarding the sample is missing. Since our network involves temporal correlations between multiple windows, it manages to output a more complete description of the entire structure. Moreover, our reconstructions were more consistent over time, namely, the predicted structure did not change significantly between frames in comparison to the results of blind inpainting. Notably, the network was capable to reconstruct random movements of filaments, despite the fact it was trained on rotations and shifts only. This suggests that the network training was general enough to avoid overfitting to simulations.

As in all model-based neural-net reconstruction algorithms, the network’s ability to generalize will always be limited by the training data, and caution should be exercised when applying the method; specifically, training data must resemble the experimental structure to avoid hallucinations^25–27^ To validate the applicability of DBlink to structures with higher structural complexity than filaments, we tracked mitochondrial dynamics. First, we trained the neural network on simulated mitochondria-like structures, drifting and wobbling in time. Then, we used weak-affinity, non-covalent fluorophore labels that allow extended observation time^28^. The combination of these dyes with our high spatiotemporal reconstruction, enables tracking dynamics in live cells at high spatiotemporal resolution in SMLM imaging over long observation times.

Future work can include extensions to other structures in live cells, e.g.: endoplasmic reticulum (ER), vimentin filaments, etc., expanding the types of motion in the training data to simulate elongation, contraction, wobbling and more complex dynamics, and systematic parameter optimization, e.g. optimal sample densities, sample-motion-rate to acquisition-rate ratio, and more. Our neural network based spatiotemporal interpolation could enable higher quality observation and ultimately facilitate discovery in various applications including cellular dynamics^29^, colocalization of nano-particles with organelles^30,31^, synthetic materials^32^, and more.

## Methods

### Training scheme

To train the neural network we used 1000 pairs of localization videos and ground truth structure videos (see SI for info) as a training set, and 250 pairs as validation set. Our loss function is comprised of three main terms: (i) Mean Squared Error (MSE); (ii) consistency loss; (iii) total variation loss. The consistency loss is calculated by the sum of pixelwise distance between every two adjacent frames

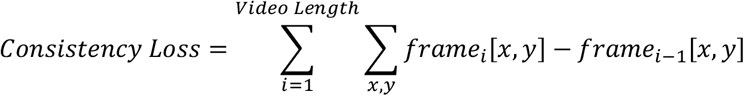

Additionally, we used the Adam optimizer (betas = [0.99, 0.999]) with reduce on plateau mechanism (patience = 3). The network was trained on a single Titan RTX GPU for approximately 3 days.

### Dynamic data acquisition

#### Drifting microtubules

To prepare cells for imaging, we cleaned cover glasses (#1.5H, 22×22 mm, Marienfeld) in an ultrasonic bath (DCG-120H, mrc) with 5% Decon90 at 60 °C for 30 min. Then we washed the cover glasses with water, incubated them in ethanol absolute for 30 min, and sterilized them with 70% filtered ethanol for 30 min. The slides were then seeded with COS7 cells and grown for 24 h in a six-well plate using phenol-free Dulbecco′s Modified Eagle′s medium (Gibco) with 1 g/l D-Glucose (i.e., low glucose), supplemented with fetal bovine serum (Biological industries), penicillin–streptomycin and glutamine at 37 °C and 5% CO2. Cells were fixed with 4% paraformaldehyde in PBS (37 °C, pH 7.3) for 20 min, washed, and incubated in 0.3 M glycine/PBS solution for 15 min. The cover glasses were transferred into a clean six-well plate, incubated for 10 min in permeabilization solution (0.25% Triton-x in PBS), washed 3 times with PBS and incubated in a blocking solution for 2 h (10% goat serum, 3% BSA, 2.2% glycine in PBS). The cells were then immunostained overnight with anti-α-tubulin-AF647 (ab190573, Abcam) diluted 1:500 in the blocking buffer. After staining, the samples were washed five times with PBS. To prevent detachment of the anti-tubulin antibodies, the sample was again treated with 4% paraformaldehyde in PBS (pH 7.3) for 10 min, washed, and incubated in 0.3 M glycine/PBS solution for 10 min.

For super-resolution imaging, a PDMS chamber was attached to a glass coverslip holding fluorescently labeled COS7 cells. Blinking buffer^33^ (Smart Kit - Super resolution buffer, Abbelight) was added and a clean coverslip was placed on top while minimizing any residual air bubbles in the chamber. The sample was illuminated with 640 nm laser (∼ 300 W/cm^2^), and 10,000 images with 50 ms exposure time were acquired (Photometrics Prime 95B).

### Cell Culture – mitochondria experiment

U2OS cells were cultured in T-75 flasks (Greiner) at 37? and 5% CO2 in Dulbecco’s Modified Eagle Medium (DMEM) / F-12 (Gibco, Thermo Fisher, USA) containing 10% (v/v) fetal bovine serum (FBS) (Corning, USA), 1% penicillin-streptomycin (w/v) (Gibco, ThermoFisher, USA) and 1% GlutaMAX (v/v) (Gibco, USA).

Cells were transiently transfected with the plasmid pCDNA5/FRT/TO-COX8A-HaloTag7. For this purpose, 2 × 10^4^ U2OS cells were seeded on fibronectin-coated 8-well chamber (Sarstedt, Germany). After 24 h incubation (37?, 5% CO_2_), cells were transfected using Lipofectamine 3000 transfection reagent (Gibco, Thermo Fisher, USA). Briefly, 0.31 μL Lipofectamine 3000 was diluted in 10.42 μL OptiMEM medium (Gibco, Thermo Fisher, USA), and 210 ng vector DNA was diluted in 10.42 μL OptiMEM medium with 0.42 μL P3000 reagent (Gibco, Thermo Fisher, USA). Diluted DNA solution was added to Lipofectamine diluent in a 1:1 ratio and incubated for 20 min at RT. After adding the DNA-lipid complex, cells were further incubated for 16-24 h at 37°C and 5% CO_2_.

Prior to imaging, the cells were washed with pre-warmed live cell imaging solution (LCIS, ThermoFisher) and temperature adjusted to avoid lateral and axial drift.

### Live cell imaging

Live cell data of microtubules was generously provided as reported previously in R. Tachibana et al^24^. Briefly, SMLM imaging was carried out using an inverted fluorescence microscope (Eclipse Ti-E; Nikon) with an oil-immersion objective (CFI Apo TIRF 100X Oil, NA 1.49; Nikon), and irradiation laser at wavelength of 561 nm (Sapphire 561 LP; Coherent). The microtubules were labeled by 4(5)-Halo-HMCR550 conjugated to HaloTag proteins. For more details on sample preparation see R. Tachibana et al^24^. For live-cell confocal imaging of U2OS expressing COX8-HaloTag7, the HT ligand JK114/HSAm carrying the fluorophore JF635 was added to the live cell imaging solution (LCIS, ThermoFisher, USA) at a final concentration of 500 nM. After an incubation time of 10 minutes, confocal microscopy was carried out on a Leica SP8 (Leica, Germany) equipped with an oil immersion objective (HC PL APO CS2 63x, NA 1,4) and an 633nm HeNe laser. Fluorophores were excited with an intensity setting of 1% 633 nm and a pinhole diameter of 1 Airy Unit. 300 frames were acquired using the HyD detector with a gain of 100 and at a scan speed of 400 Hz in xyt acquisition mode. Leica LASX software was used for the microscope control and data acquisition.

For live-cell SMLM imaging of U2OS expressing COX8-HaloTag7, the HT ligand JK114/HSAm carrying the fluorophore JF635 was added to the pre-warmed live cell imaging solution (LCIS, ThermoFisher, USA) at a final concentration of 1 nM. After an incubation time of 10 minutes, imaging was carried out on a N-STORM microscope (Nikon, Japan) equipped with an oil immersion objective (Apo, 100x, NA 1.49) and an EMCCD camera (DU-897U-CS0-#BV, Andor Technology, Ireland). Fluorophores were excited with a collimated 647 nm laser beam at an intensity of 0.4 kW/cm2 (measured at the objective back focal plane) at highly inclined and laminated optical sheet (HILO) mode. 20,000-60,000 consecutive frames were acquired at 50 Hz in active frame transfer mode with an EMCCD gain of 200, a pre amp gain of 3, readout mode of 17MHz and at an effective pixel size of 158 nm. NIS Elements (Nikon, Japan), LCControl (Agilent, USA), and Micro-Manager were used for the optical setup and the data acquisition.

## Code availability

The code is available online at: https://github.com/alonsaguy/DBlink.

## Acknowledgements

Live cell data of dynamic microtubules was generously shared with us by Prof. Yasuteru Urano and Prof. Mako Kamiya of The University of Tokyo. We thank Julian Lau and Kai Johnsson (MPI Medical Research, Heidelberg, and EPFL) for kindly providing the plasmid COX8A-HaloTag7. M.H. and S.J. gratefully acknowledge funding by LOEWE (FCI) and the Deutsche Forschungsgemeinschaft (DFG, German Research Foundation) – SFB1177; INST 161/926-1 FUGG. A.S. has received funding from the European Union’s Horizon 2020 research and innovation program under grant agreement No. 802567 -ERC-Five-Dimensional Localization Microscopy for Sub-Cellular Dynamics. Y.S. is supported by the Zuckerman Foundation.

## References

1. Hell, S. W. & Wichmann, J. Breaking the Diffraction Resolution Limit By Stimulated-Emission - Stimulated-Emission-Depletion Fluorescence Microscopy. Opt. Lett. 19, 780–782 (1994).

2. Gustafsson, M. G. L. Surpassing the lateral resolution limit by a factor of two using structured illumination microscopy. J. Microsc. 198, 82–87 (2000).

3. Betzig, E. et al. Imaging intracellular fluorescent proteins at nanometer resolution. Science. 313, 1642–1645 (2006).

4. Rust, M. J., Bates, M. & Zhuang, X. Sub-diffraction-limit imaging by stochastic optical reconstruction microscopy (STORM). Nat. Methods 3, 793–796 (2006).

5. Schnitzbauer, J., Strauss, M. T., Schlichthaerle, T., Schueder, F. & Jungmann, R. Super-resolution microscopy with DNA-PAINT. Nat. Protoc. 12, 1198–1228 (2017).

6. Alexey, S. & M., H. R. Wide-field subdiffraction imaging by accumulated binding of diffusing probes. Proc. Natl. Acad. Sci. 103, 18911–18916 (2006).

7. Ouyang, W., Aristov, A., Lelek, M., Hao, X. & Zimmer, C. Deep learning massively accelerates super-resolution localization microscopy. Nat. Biotechnol. 36, 460–468 (2018).

8. Wang, Y. et al. Blind sparse inpainting reveals cytoskeletal filaments with sub-Nyquist localization. Optica 4, 1277–1284 (2017).

9. Nehme, E., Weiss, L. E., Michaeli, T. & Shechtman, Y. Deep-STORM: super-resolution single-molecule microscopy by deep learning. Optica 5, 458 (2018).

10. Nehme, E. et al. DeepSTORM3D: dense 3D localization microscopy and PSF design by deep learning. Nat. Methods 17, 734–740 (2020).

11. Speiser, A. et al. Deep learning enables fast and dense single-molecule localization with high accuracy. Nat. Methods 18, 1082–1090 (2021).

12. Wu, Y. & Shroff, H. Faster, sharper, and deeper: structured illumination microscopy for biological imaging. Nature Methods (2018).

13. Priessner, M. et al. Content-aware frame interpolation (CAFI): Deep Learning-based temporal super-resolution for fast bioimaging. bioRxiv 2021.11.02.466664 (2021).

14. Chen, R. et al. Deep-Learning Super-Resolution Microscopy Reveals Nanometer-Scale Intracellular Dynamics at the Millisecond Temporal Resolution. bioRxiv 2021.10.08.463746 (2021).

15. Nehme, E., Weiss, L. E., Michaeli, T. & Shechtman, Y. Deep-STORM: super-resolution single-molecule microscopy by deep learning. Optica 5, 458 (2018).

16. Ovesný, M., KříŽek, P., Borkovec, J., Svindrych, Z. & Hagen, G. M. ThunderSTORM: a comprehensive ImageJ plug-in for PALM and STORM data analysis and super-resolution imaging. Bioinformatics 30, 2389–90 (2014).

17. Su, Y.-T., Lu, Y., Chen, M. & Liu, A.-A. Spatiotemporal Joint Mitosis Detection Using CNN-LSTM Network in Time-Lapse Phase Contrast Microscopy Images. IEEE Access 5, 18033–18041 (2017).

18. Yu, Y., Si, X., Hu, C. & Zhang, J. A Review of Recurrent Neural Networks: LSTM Cells and Network Architectures. Neural Comput. 31, 1235–1270 (2019).

19. Shariff, A., Murphy, R. F. & Rohde, G. K. A generative model of microtubule distributions, and indirect estimation of its parameters from fluorescence microscopy images. Cytom. Part A 77, 457–466 (2010).

20. Banterle, N., Bui, K. H., Lemke, E. A. & Beck, M. Fourier ring correlation as a resolution criterion for super-resolution microscopy. J. Struct. Biol. 183, 363–367 (2013).

21. Descloux, A., Grußmayer, K. S. & Radenovic, A. Parameter-free image resolution estimation based on decorrelation analysis. Nat. Methods 16, 918–924 (2019).

22. Kompa, J. et al. Exchangeable HaloTag Ligands (xHTLs) for multi-modal super-resolution fluorescence microscopy. bioRxiv 2022.06.20.496706 (2022).

23. Lefebvre, A. E. Y. T., Ma, D., Kessenbrock, K., Lawson, D. A. & Digman, M. A. Automated segmentation and tracking of mitochondria in live-cell time-lapse images. Nat. Methods 18, 1091–1102 (2021).

24. Tachibana, R. et al. Design of spontaneously blinking fluorophores for live-cell super-resolution imaging based on quantum-chemical calculations. Chem. Commun. 56, 13173–13176 (2020).

25. Möckl, L., Roy, A. R. & Moerner, W. E. Deep learning in single-molecule microscopy: fundamentals, caveats, and recent developments [Invited]. Biomed. Opt. Express 11, 1633 (2020).

26. Matlock, A. & Tian, L. Physical model simulator-trained neural network for computational 3d phase imaging of multiple-scattering samples. arXiv Prepr. arXiv2103.15795 (2021).

27. Belthangady, C. & Royer, L. A. Applications, promises, and pitfalls of deep learning for fluorescence image reconstruction. Nat. Methods (2019).

28. Spahn, C., Grimm, J. B., Lavis, L. D., Lampe, M. & Heilemann, M. Whole-Cell, 3D, and Multicolor STED Imaging with Exchangeable Fluorophores. Nano Lett. 19, 500–505 (2019).

29. Wensel, T. G., Potter, V. L., Moye, A., Zhang, Z. & Robichaux, M. A. Structure and dynamics of photoreceptor sensory cilia. Pflügers Arch. - Eur. J. Physiol. 473, 1517–1537 (2021).

30. Guggenheim, E. J. et al. Comparison of confocal and super-resolution reflectance imaging of metal oxide nanoparticles. PLoS One 11, e0159980 (2016).

31. van der Zwaag, D. et al. Super Resolution Imaging of Nanoparticles Cellular Uptake and Trafficking. ACS Appl. Mater. Interfaces 8, 6391–6399 (2016).

32. Pujals, S., Feiner-Gracia, N., Delcanale, P., Voets, I. & Albertazzi, L. Super-resolution microscopy as a powerful tool to study complex synthetic materials. Nat. Rev. Chem. 3, 68–84 (2019).

33. Nahidiazar, L., Agronskaia, A. V., Broertjes, J., van den Broek, B. & Jalink, K. Optimizing Imaging Conditions for Demanding Multi-Color Super Resolution Localization Microscopy. PLoS One 11, e0158884 (2016).

