## Supplementary Information for "DBlink: Dynamic localization microscopy in super spatiotemporal resolution via deep learning"

### Filament simulation

The initial ground truth structures were simulated according to the model suggested by Shariff, et al[1]. Then, we applied affine temporally changing transformations to the original structure over predetermined video length (see Table S1). We applied on the initial ground truth structure two types of movements: global shift and global rotation. The shift velocities were chosen from a uniform distribution in the range of [-4, 4] nm per frame, and the rotation velocities were chosen from a uniform distribution in the range of [-3, 3] degrees per frame. Next, we randomly chose the number of blinking events per frame according to a blinking density parameter that states the percentage of the structure that would blink at each frame. We determined the position of each simulated blinking event according to the ground truth structure with additional localization noise randomly chosen from a uniform distribution in the range of [-20, 20] nm. Finally, we added additional localizations at random positions in the field of view (FOV) as noise. The result was pairs of simulated localization video and underlying dynamic structure video.

| Parameters | Video length [frames] | Pixel size [um] | Field of view [um] | Blink density [%] |
|---|---|---|---|---|
| Values | 1000 | 0.16 | 5.12 x 5.12 | 0.2 |

Table S1: Simulation parameters.

### Mitochondria simulation

Here we followed a similar scheme of simulation, but we changed the ground truth and the simulation parameters. First, we chosen N random center-of-mass (CM) positions for N mitochondria in the simulated field of view (FOV). Then, around each position we have chosen a random number of edge points from a uniform distribution of [30, 50] points. Each point was assigned with an angle in the range of [0, $2\pi$] and a distance from the center of mass according to the known size of mitochondria (see Table S2). Finally, we acquired the ground truth structures of each mitochondrion by drawing a polygon based on the randomly chosen edge points.

| Parameters | Video length [frames] | Pixel size [um] | Field of view [um] | Distance from CM [um] | Blink density [%] |
|---|---|---|---|---|---|
| Values | 1000 | 0.16 | 5.12 x 5.12 | 0.5 – 1.2 | 0.5 |

Table S2: Simulation parameters.

We have simulated two types of dynamic movements for each mitochondrion: global shift, with velocities in the range of [-20, 20] nm per frame; and mitochondrion warping. The warping was done by choosing K edge points and move them periodically according to a sine function.

The blinking videos were simulated in a similar fashion to the simulations of filaments, but some parameters have changed (see Table S2).

### Neural network architecture

Super spatio-temporal resolution reconstruction falls within the domain of sequence-to-sequence (seq2seq) objectives. In our case, the input is a sequence of high-precision localization maps of single molecules in an SMLM experiment, and the output is a sequence of images containing high-resolution reconstruction of the imaged structure.

Previous work has shown that combining information from multiple frames is beneficial in means of reconstruction accuracy and temporal resolution improvement[2,3]. However, the suggested algorithms are based on CNNs which are sub-optimal solution for seq2seq objectives. A more commonly used architecture for seq2seq tasks is Recurrent neural network (RNN).

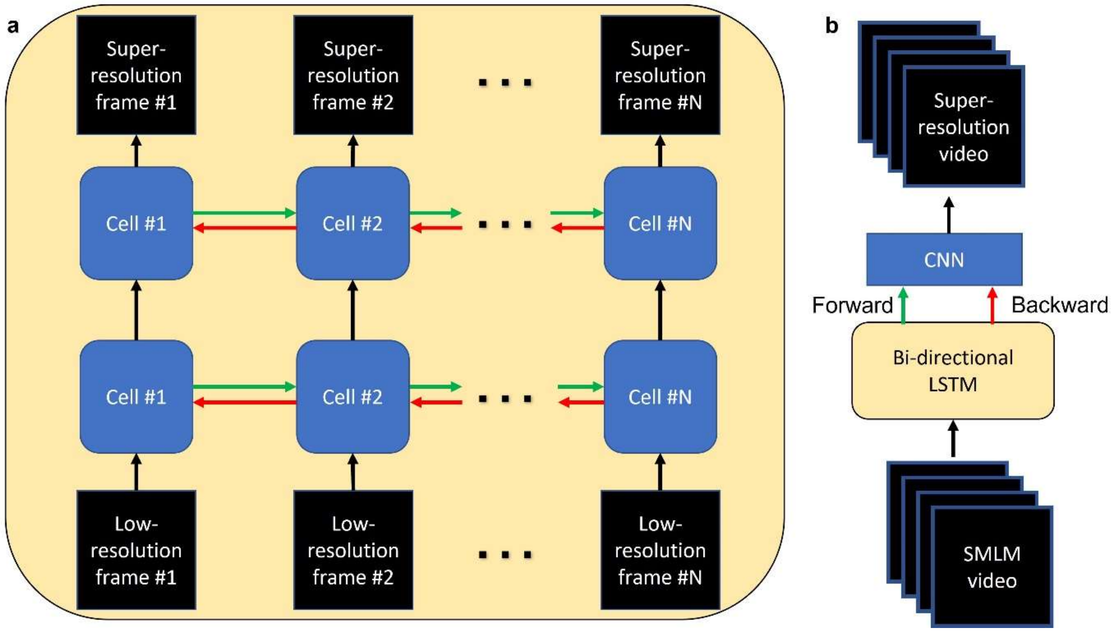


Figure S1: Neural network architecture. **a** We implemented bi-directional LSTM network with two layers. The first layer blocks get as input single low-resolution frames and the hidden states of the previous block. The second layer blocks output single super-resolved frames. In the forward pass (green arrows) the information propagates chronologically, while in the backward pass (red arrows) the images are inserted to the same network in reverse order. **b** The output frames of both the forward and the backward pass are inserted to a CNN as two input channels. The output of the CNN is the super-resolved reconstruction of the entire video.

RNNs combine temporal information along the input sequence to provide better reconstructions in the output sequence. The weights in each RNN block are recycled during the inference process; therefore, RNNs are composed of less parameters than CNNs. Nevertheless, RNNs outperforms CNNs in many seq2seq objectives. In our work, we implement a variant of RNNs named bi-directional long short term memory (LSTM) network (Supplementary Figure S1). LSTMs are known for their ability to propagate important information throughout long input sequences. This advantage, along with the low memory demand, makes them perfect for the analysis of videos.

In addition to the suggested architecture, we have added another post-processing step to our analysis. In this step, we transform the output image to binary mask by defining all the pixels with values greater than some threshold as ones and the rest as zeros. Since the output image of the neural network $I(x, y)$ may be seen as a heatmap indicating the confidence of the network in the presence of a structure in each reconstructed pixel, we weighted each pixel in the binary map $B(x, y)$ according to the network confidence. Therefore, we drew a patch around each pixel and multiplied this patch by a 2D gaussian with standard deviation that equals to one over the original pixel value:

$$B(x_i, y_i) = \begin{matrix} 1\,, & I(x_i, y_i) > threshold \\ 0\,, & I(x_i, y_i) < threshold \end{matrix}$$

$$Final\ output(x, y) = B(x, y) \cdot \frac{1}{\sqrt{2\pi \cdot I(x_i, y_i)}} \cdot e^{\frac{-(x-x_i)^2-(y-y_i)^2}{2\cdot I^2(x_i,y_i)}} \quad , x, y \in \{patch\ around\ x_i, y_i\}$$

This function would decrease the pixel intensity where the network confidence is low and maintain high pixel intensity otherwise. The resulting frame of this analysis would keep the intensity of high confidence pixels and reduce the intensity of low confidence pixels.

### Reconstruction accuracy quantification

The ground truth in our simulations were binary masks containing ones on the sample localization and zeros on the background. In the case of experimental data, we either did not possess any information regarding the ground truth structure (in the case of live cell imaging) or we possessed the reconstruction of the data based on Deep-STORM's predictions. The outputs of our network were heatmaps containing different values in the range of [0, 1]. Higher pixel values meant that the network had higher confidence in estimating the structure at those pixels.

Finding the optimal accuracy measure for comparison between the predictions and ground truth is not a trivial task. We have considered several accuracy measures for the quantification of our reconstruction performance. The pixel-wise mean squared error (MSE) is a widely used measure for this purpose; however, in some cases it poorly describes the quality as we would expect. For example, when the sample is small relative to the FOV, most of the pixels in the ground truth image would have zero intensity. Therefore, consistently predicting matrices full of zero values would yield a very low error using the MSE. Structure similarity (SSIM)[4] will suffer from similar problems as MSE, due to the fact it relies on comparison between the mean intensity and standard deviation of the predicted image and the ground truth. Jaccard index[5] might be used to describe the similarity between two groups: the group of predicted localizations and the group of ground truth localizations. But in our case, we compare matrices and not localization lists and it is hard to compare between the predicted heatmaps provided by our neural network and the ground truth binary maps representing the sample.

Therefore, we have decided to quantify DBlink performance on simulated data according to two different quantities: the reconstruction fidelity to the ground truth structure; and the network hallucinations displayed in its reconstructions. The reconstruction fidelity term is measured by the following steps: binarizing the predicted image based a predefined threshold of half the maximal intensity; counting the number of pixels marked as ones in both the predicted image and the ground truth; dividing that number by the total number of pixels marked as ones in the ground truth image. The hallucination term was measured by the following steps: summing the number of pixels marked as ones in the predicted image and as zeros at the ground truth image; dividing this number by the number of pixels marked as zeros in the ground truth.

In the experiment that contained unwanted stage drift, we quantified the accuracy as follows: First, we generated the ground truth image using ThunderSTORM reconstruction with drift correction and density filter tools. Next, we shifted back our reconstructed video frames according to the framewise drift prediction. Then, we binarized both our reconstructions and the ground truth reconstruction with thresholds that equal to the 75$^{th}$ percentile of each image intensity histogram (Supplementary Fig. S2). Finally, we quantified the reconstruction accuracy by measuring the cross-correlation between the re-shifted reconstructions $(\hat{y}_i)$ and the static ground truth image $(y)$. The final normalized term we used is:

$$Accuracy_i = \frac{\max(\hat{y}_i \star y)}{\sqrt{(\hat{y}_i \star \hat{y}_i) \cdot (y \star y)}}$$

Where $\star$ marks the cross-correlation operator. The mean accuracy we obtained was 0.89.

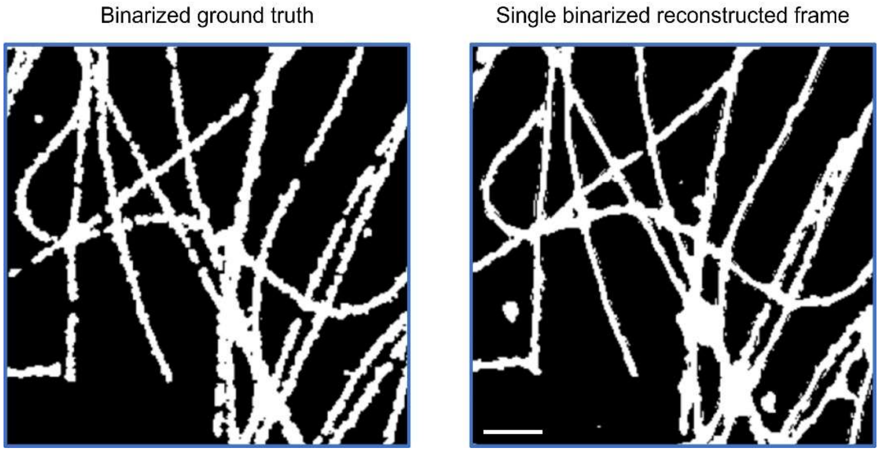


Figure S2: Quantifying the reconstruction accuracy in drifting sample experiment. Left: The ground truth structure obtained by ThunderSTORM reconstruction in addition to application of drift correction and density filtration. Right: A single reconstructed frame of DBlink. Both images were binarized according to the 75th percentile of each image. Scale bar = 2 $\mu m$.

In the experiment that contained camera rotation, due to the finite numerical limitation to exactly rotate and shift back each frame we quantified a different property of our reconstructions – the consistency. To do so, we have measured the cross-correlation between every two frames in the reconstructed video:

$$Accuracy_{ij} = \frac{\max(\hat{y}_i \star \hat{y}_j)}{\sqrt{(\hat{y}_i \star \hat{y}_i) \cdot (\hat{y}_j \star \hat{y}_j)}}$$

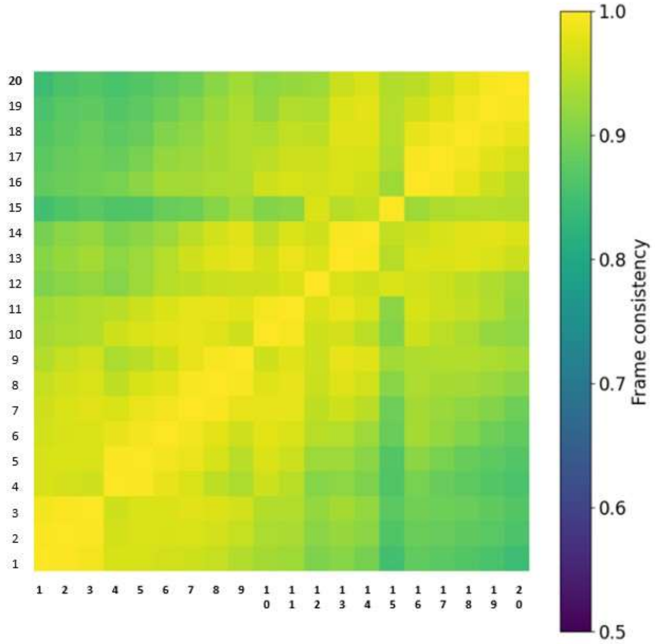


Figure S3: Consistency quantification. We measured the normalized cross-correlation between every two frames in the reconstructed video. The diagonal values mark the autocorrelations of each frame with itself; hence, they contain ones.

The result of this measurement is a matrix containing ones in the diagonal and normalized cross-correlations elsewhere. We achieved a mean consistency term of 0.91, over 20 neighboring frames, indicating that our reconstructed structure does not change throughout the reconstructed video (Supplementary Fig. S3).

### Spatial resolution quantification

We have quantified the spatial resolution according to Fourier ring correlation (FRC)[6]. In this method, we used DBlink reconstruction of static data along with super-resolution reconstruction of the same structure using ThunderSTORM. Then, we multiplied the Fourier transform of each subsample. Finally, we measure the mean value of the multiplication image over rings with an increasing size. When the mean pixel intensity of a ring drops below a certain threshold, we mark the radius of that ring as the maximal spatial frequency that occurs in our reconstruction (Supplementary Fig. S4). The resolution is estimated by the dividing one by the maximal spatial frequency we achieved. We used the previously suggested $2\sigma$ threshold as our decision threshold. This threshold is computed by dividing 2 over the square root of half the number of pixels in each ring.

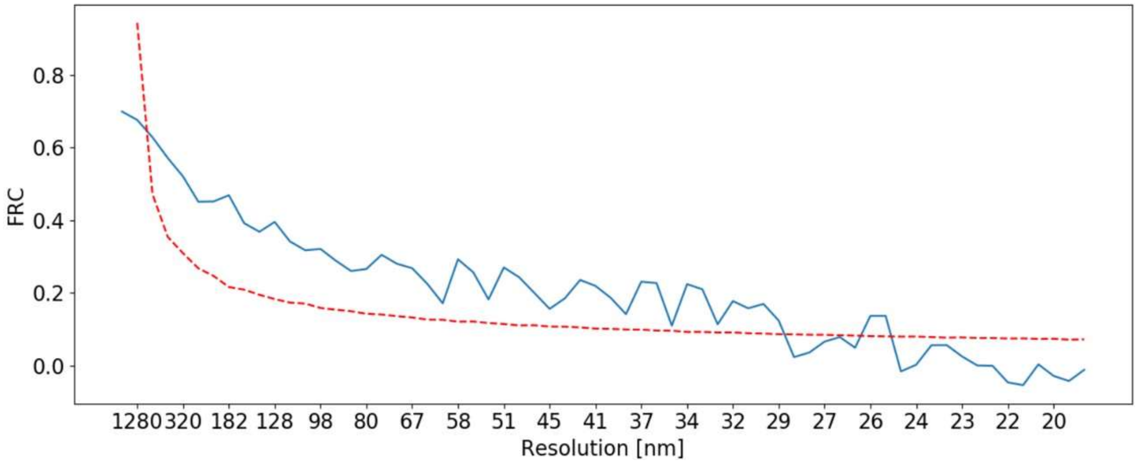


Figure S4: Fourier ring correlation analysis for spatial resolution quantification. First, we take two random subsamples of the data; then, we multiply the Fourier transform of the subsamples. Finally, we calculate the FRC according to the mean intensity value of all the pixels in a ring increasing in size. The resolution is determined according one over the cut-off frequency we achieved in the meeting point between the FRC curve and the predetermined threshold.

In addition to FRC, we have used another previously algorithm for resolution estimation, decorrelation analysis[7]. According to decorrelation analysis DBlink obtained spatial resolution of 37 nm (Supplementary Fig. S5).

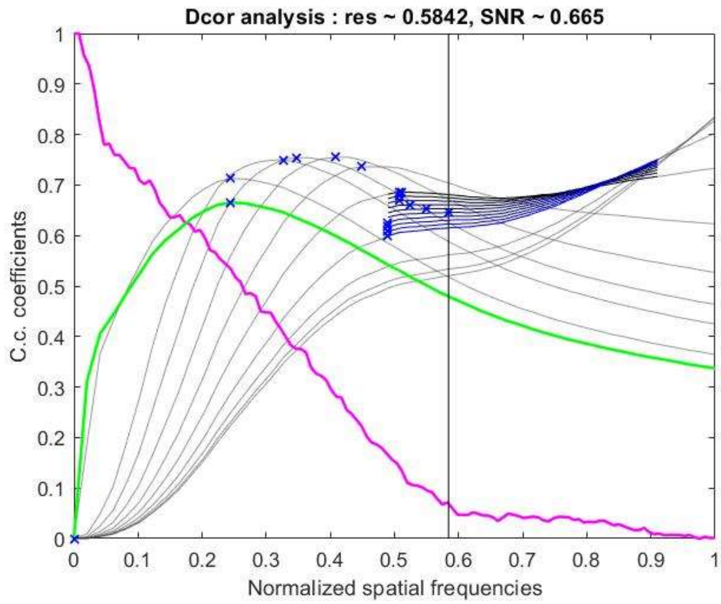


Figure S5: Decorrelation analysis. The output of decorrelation analysis algorithm published by A. Descloux et al[7]. The maximal spatial frequency in our reconstruction was the ~58th percentile of the maximal achievable frequency in our system. In our experiment this number matched spatial resolution of ~35 nm.

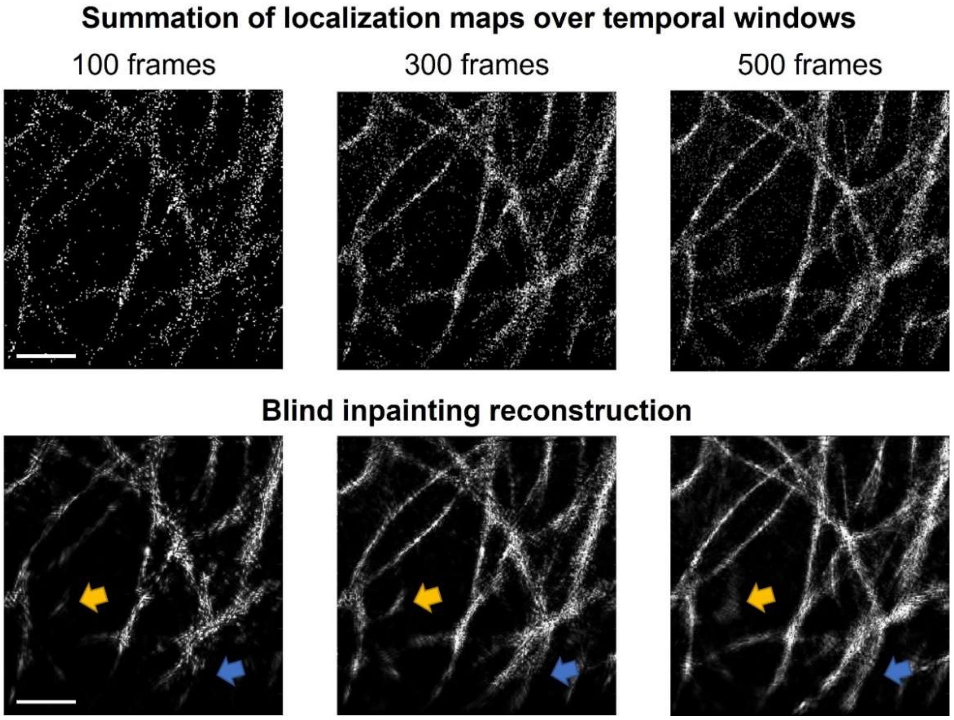


Figure S6: Blind inpainting evaluation. Upper row: summation frame of localization maps over temporal windows with varying length. Bottom row: reconstruction of the summation frame by applying blind inpainting algorithm on it. All the reconstructions in the bottom row managed to filter noise and emphasize relevant features of the sample. However, summing 100 localization frames is not sufficient for the reconstruction of the entire sample; on the other hand, summing 500 frames generated motion blur that blind inpainting could not resolve (yellow and blue arrows). Empirically, the best compromise between motion blur and reconstruction accuracy was obtained when we summed 300 frames. Scale bar = 2.5 um.

### References


1. Shariff, A., Murphy, R. F. & Rohde, G. K. A generative model of microtubule distributions, and indirect estimation of its parameters from fluorescence microscopy images. *Cytom. Part A* **77**, 457–466 (2010).
2. Ouyang, W., Aristov, A., Lelek, M., Hao, X. & Zimmer, C. Deep learning massively accelerates super-resolution localization microscopy. *Nat. Biotechnol.* **36**, 460–468 (2018).
3. Nehme, E., Weiss, L. E., Michaeli, T. & Shechtman, Y. Deep-STORM: super-resolution single-molecule microscopy by deep learning. *Optica* **5**, 458 (2018).
4. Wang, Z., Bovik, A. C., Sheikh, H. R. & Simoncelli, E. P. Image quality assessment: from error visibility to structural similarity. *IEEE Trans. Image Process.* **13**, 600–612 (2004).
5. Jaccard, P. THE DISTRIBUTION OF THE FLORA IN THE ALPINE ZONE.1. *New Phytol.* **11**, 37–50 (1912).
6. Banterle, N., Bui, K. H., Lemke, E. A. & Beck, M. Fourier ring correlation as a resolution criterion for super-resolution microscopy. *J. Struct. Biol.* **183**, 363–367 (2013).
7. Descloux, A., Grußmayer, K. S. & Radenovic, A. Parameter-free image resolution estimation based on decorrelation analysis. *Nat. Methods* **16**, 918–924 (2019).
